# Targeted Nanopore Sequencing with Cas9 for studies of methylation, structural variants, and mutations

**DOI:** 10.1101/604173

**Authors:** Timothy Gilpatrick, Isac Lee, James E. Graham, Etienne Raimondeau, Rebecca Bowen, Andrew Heron, Fritz J Sedlazeck, Winston Timp

## Abstract

Nanopore sequencing technology can rapidly and directly interrogate native DNA molecules. Often we are interested only in interrogating specific areas at high depth, but conventional enrichment methods have thus far proved unsuitable for long reads^1^. Existing strategies are currently limited by high input DNA requirements, low yield, short (<5kb) reads, time-intensive protocols, and/or amplification or cloning (losing base modification information). In this paper, we describe a technique utilizing the ability of Cas9 to introduce cuts at specific locations and ligating nanopore sequencing adaptors directly to those sites, a method we term ‘nanopore Cas9 Targeted-Sequencing’ (nCATS).

We have demonstrated this using an Oxford Nanopore MinION flow cell (Capacity >10Gb+) to generate a median 165X coverage at 10 genomic loci with a median length of 18kb, representing a several hundred-fold improvement over the 2-3X coverage achieved without enrichment. We performed a pilot run on the smaller Flongle flow cell (Capacity ~1Gb), generating a median coverage of 30X at 11 genomic loci with a median length of 18kb. Using panels of guide RNAs, we show that the high coverage data from this method enables us to (1) profile DNA methylation patterns at cancer driver genes, (2) detect structural variations at known hot spots, and (3) survey for the presence of single nucleotide mutations. Together, this provides a low-cost method that can be applied even in low resource settings to directly examine cellular DNA. This technique has extensive clinical applications for assessing medically relevant genes and has the versatility to be a rapid and comprehensive diagnostic tool. We demonstrate applications of this technique by examining the well-characterized GM12878 cell line as well as three breast cell lines (MCF-10A, MCF-7, MDA-MB-231) with varying tumorigenic potential as a model for cancer.

**Contributions:** TG and WT constructed the study. TG performed the experiments. TG, IL, and FS analyzed the data. TG, JG, ER, RB and AH and developed the method. TG and WT wrote the paper

## Introduction

Nanopore sequencing operates by reading the DNA base sequence through fluctuations in ionic current as a DNA molecule threads through a protein pore embedded in a membrane. This has been commercialized by Oxford Nanopore; their flagship product is the minION flow cell, which generates 10+ gigabases of data. There is a need for tests that can affordably scale to the demands of diagnostic laboratories to enable fast diagnostics with high sequencing depth at clinically informative loci. However, current capture assays are still logistically challenging for long-read sequencing. Existing capture methods adapted to long read sequencing include CATCH-seq (dual cuts with Cas9 followed by size selection)^2^, ligation to cuts with a less permissive endonuclease^3^, cloning the region into an expression plasmid^4^, region amplification^5^, and long fragment hybridization capture methods^1,6^. These methods are limited by the loss of native modifications, short read length, high input, low yield, or long protocols (>18hrs). We describe here an enrichment strategy using targeted cleavage with Cas9 for ligating nanopore sequencing adaptors (nanopore Cas9-Targeted Sequencing, or ‘nCATS’) and show the use of this method for simultaneously assessing single nucleotide variants (SNVs), structural variants (SVs) and CpG methylation. The method can be completed on a benchtop in several hours, needs only ~3ug of genomic DNA and can be easily adapted to target a large number of loci in a single reaction. The potential of this technique, combined with the low capital investment required for nanopore sequencing instruments (~$1K), could put a targeted sequencing assay in the hands of every pathology department, to evaluate cellular DNA for CpG methylation, structural rearrangements, and survey for nucleotide mutations.

## Results

We enrich by selective ligation of nanopore sequencing adaptors at fresh cut sites created by active Cas9. To achieve enrichment, all preexisting DNA ends are dephosphorylated prior to the addition of the Cas9/guide RNA ribonucleoprotein complex (RNP). Cuts introduced by the RNPs thereby represent the majority of DNA ends with a 5-phosphate group, and as a result nanopore sequencing adaptors are preferentially ligated to the DNA ends made by Cas9 cleavage (**Figure 1A**). By sequencing the native DNA strands, we avoid the need for PCR amplification and maintain any modified nucleotides; nanopore sequencing can distinguish modifications using raw electrical data^7^. Using human breast cell lines, we demonstrate applications of this method for characterizing breast cancer, focusing on genes where transcription and methylation have prognostic implications as well as sites believed to harbor large (>1kb) chromosomal deletions and finally single nucleotide variants (SNVs). We used the well-characterized GM12878 cell line to validate our ability to examine these features with the deeper coverage data from the nCATS method through comparison to annotated variants^8,9^.

**Figure 1:**
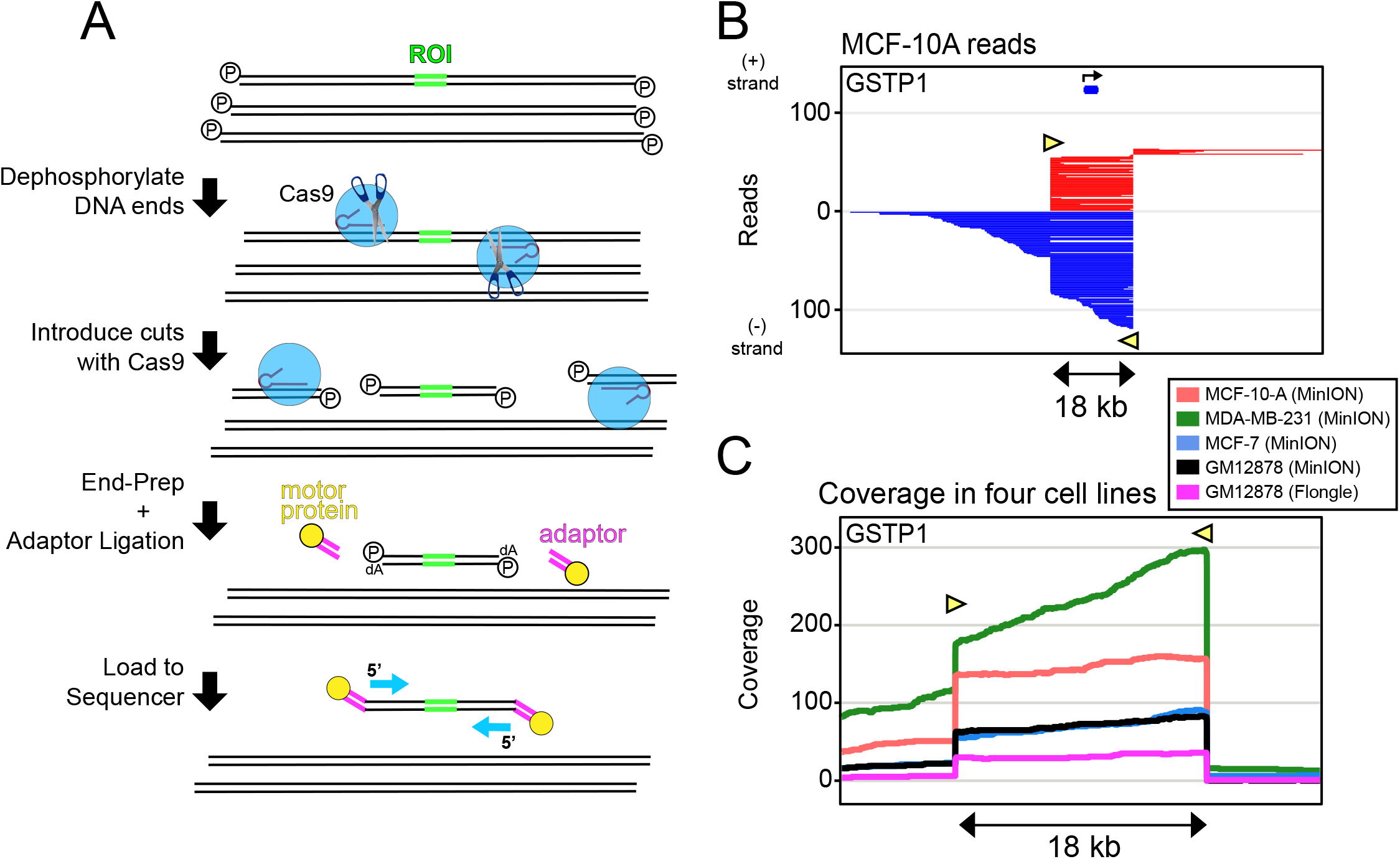
Method schematic and coverage data. (A) Schematic of Cas9 enrichment operation. ROI = region of interest. DNA ends are first dephosphorylated, new cuts introduced with Cas9/guideRNA complex, nanopore sequencing adaptors are ligated to cuts around the ROI and and the sample is loaded to the nanopore sequencer. (B) Representative read-plots for the MCF-10-A cell line at GSTP1. Yellow triangles show Cas9 cut site and guideRNA direction (C) Coverage plots at the GSTP1 gene in the three breast cell lines and GM12878 cell line

At each site evaluated, we used Cas9 to introduce two cuts flanking the region of interest. For sites without structural variations, the guide RNAs (gRNAs) were placed 20-30kb apart, and for sites with deletions, the gRNAs were designed to flank breakpoints by ~5kb. We targeted 10 loci (**Supplementary Table 1**) in four cell lines: three breast cell lines (MCF-10A, MCF-7, and MDA-MB-231) and the well-characterized GM12878 lymphoblast cell line. For studying DNA methylation and structural variants, regions of interest were selected based on previous whole-genome nanopore data from our lab^10^ as well as existing expression data in these breast cell lines^11^. To evaluate single nucleotide mutations we selected three cancer-associated genes that have annotated mutations in the MDA-MB-231 cell line^12^.

### Yield and Coverage

Starting with ~3μg of genomic DNA from each cell line, we captured 10 loci, dedicating a single MinION flow cell to each sample, which resulted in a local coverage ranging from 20X to 800X. We repeated the capture experiment in GM12878 using the newer, smaller flongle flow cell, which generated a local coverage from 10X to 70X. For the flongle sequencing run, we added guideRNAs flanking the BRCA1 locus, but the size of this gene (>80kb) is larger than we were conventionally targeting with this approach and resulted in a low average coverage of only 3X. The total yield per MinION flow cell ranged from 70K to 230K reads (1.7-7.6% on-target), and the flongle produced 12K reads (3.8% on-target) (Table 1). Genome-wide coverage analysis found the off-target reads to be distributed randomly across the genome, indicating they result from ligation of nanopore adaptors to random breakage points, with no clear evidence of off-target cleavage by Cas9. For example, in the GM12878 cell line, after quality filtering alignments (MAPQ > 30) there were only 2 genomic sites outside target regions where coverage reached 25X, both at repetitive peri-centromeric sites and containing reads with lower mapping quality (MAPQ 30-50), suggesting the increased coverage to be the result of alignment errors in these poorly mappable regions. In contrast, all on-target alignments had MAPQ scores >50.

**Table 1:** Average coverage and percent of total on-target reads at ten loci in four cell lines

| cell line<br>flow cell<br>total on-target reads | region<br>size<br>(kb) | GM12878 |  |  |  | MCF-10-A |  | MDA-MB-231 |  | MCF-7 |  |
| --- | --- | --- | --- | --- | --- | --- | --- | --- | --- | --- | --- |
|  |  | (MinION) |  | (Flongle) |  | (MinION) |  | (MinION) |  | (MinION) |  |
|  |  | 1388 |  | 450 |  | 4510 |  | 4192 |  | 1562 |  |
| LOCUS |  | Avg Coverage | (%) of on-target | Avg Coverage | (%) of on-target | Avg Coverage | (%) of on-target | Avg Coverage | (%) of on-target | Avg Coverage | (%) of on-target |
| GPX1 | 13.6 | 135 | 9.73% | 65 | 14.44% | 655 | 14.52% | 415 | 9.90% | 105 | 6.72% |
| GSTP1 | 17.8 | 70 | 5.04% | 30 | 6.67% | 140 | 3.10% | 225 | 5.37% | 65 | 4.16% |
| KRT19 | 18.1 | 50 | 3.60% | 20 | 4.44% | 110 | 2.44% | 250 | 5.96% | 85 | 5.44% |
| SLC12A4 | 24.4 | 175 | 12.61% | 35 | 7.78% | 585 | 12.97% | 415 | 9.90% | 330 | 21.13% |
| TPM2 | 19.6 | 110 | 7.93% | 20 | 4.44% | 335 | 7.43% | 210 | 5.01% | 65 | 4.16% |
| chr5 deletion | 18.7 | 20 | 1.44% | 10 | 2.22% | 65 | 1.44% | 140 | 3.34% | 45 | 2.88% |
| chr7 deletion | 20.0 | 75 | 5.40% | 30 | 6.67% | 340 | 7.54% | 260 | 6.20% | 80 | 5.12% |
| BRAF | 12.3 | 45 | 3.24% | 20 | 4.44% | 205 | 4.55% | 200 | 4.77% | 50 | 3.20% |
| KRAS | 16.7 | 85 | 6.12% | 25 | 5.56% | 200 | 4.43% | 250 | 5.96% | 100 | 6.40% |
| TP53 | 16.1 | 235 | 16.93% | 45 | 10.00% | 845 | 18.74% | 450 | 10.73% | 180 | 11.52% |

**Figure 1B** shows MinION reads from the MCF-10-A cell line aligning to a representative locus (*GSTP1)*. Cas9 remains bound to the upstream (5’) side of the gRNA after DNA cleavage^13^, resulting in preferential ligation of adaptors onto the 3’ side of the cut. A comparison of coverage between sequencing runs at the *GSTP1* locus is shown in **Figure 1C**. There was a difference in yield between the different guide RNAs and between cell lines, summarized in **Table 1**. This difference in coverage can be partially explained by both the stochastic formation of ribonucleoprotein complexes after combining the Cas9 with the gRNAs, as well as the different performance expected from the different gRNAs due to varying off-target mismatches and on-target binding performance. Further, a disparate number of reads were noted between cell lines which can be attributed to both (1) efforts made to keep the DNA as intact as possible, as the long strands of DNA may make it harder to get a uniform quantification; as well as (2) the aneuploidy which is known to exist in these immortalized breast cell lines. MDA-MB-231 and MCF-7 are near-triploid and hypotetraploid, with modal chromosomal numbers of 64 and 82, respectively^14,15^.

### Methylation Studies

Five of our selected loci are in the promoters of genes where expression level and promoter methylation have prognostic implications in human cancer. Sites for methylation studies were selected by examining whole-genome nanopore methylation data from these breast cell lines^10^, searching for differentially methylated promoters between the non-tumorigenic line MCF-10A and the tumorigenic lines MCF-7 and MDA-MB-231. Candidate loci were filtered for genes where methylation status is clinically informative^16–18^, and compared to existing RNA-seq expression data^11^ for differentially expressed genes.

Because nanopore sequencing directly evaluates methylation patterns on native DNA strands, we are able to observe long-range methylation information on each DNA molecule. We use read-level methylation plots to demonstrate this phased methylation information from both a MinION sequencing run and Flongle sequencing run (**Figure 2A; Supplementary Figure S1**), where we show methylation calls for evaluated CpGs directly on each read of cellular DNA. To validate the use of the nCATS method for studying DNA methylation, we compared nanopore methylation calls with existing whole genome bisulfite sequencing (WGBS) data in GM12878^19^. Using line plots to show smoothed (loess) methylation, we compare methylation patterns at each of the five regions. Specifically, we have plotted methylation data for two genes where CpG methylation is known to inform outcomes in human breast cancer (*GSTP1* and *KRT19)* (**Figure 2A**). Three additional genes (*GPX1*, *SLC12A4*, and *TPM2)* are included in **Supplementary Figure S1**. We note very similar methylation patterns between targeted nanopore and whole genome bisulfite data over all regions analyzed in GM12878, even with the noisy signal observed for many CpG sites. The aggregate correlation between the nanopore methylation calls and Illumina methylation across all five regions was 0.82 (Pearson). Dot plots comparing methylation calls at each of the five genes are included in the supplemental materials (**Supplementary Fig. S2**), wherein we observe per-CpG methylation largely clustered at points reflecting completely methylated or completely unmethylated sites. We also note that there is a substantial reduction in the noise of the methylation signature around transcriptional start sites, suggesting that CpGs which play the largest regulatory role may have less inter-cellular variation in methylation status.

**Figure 2:**
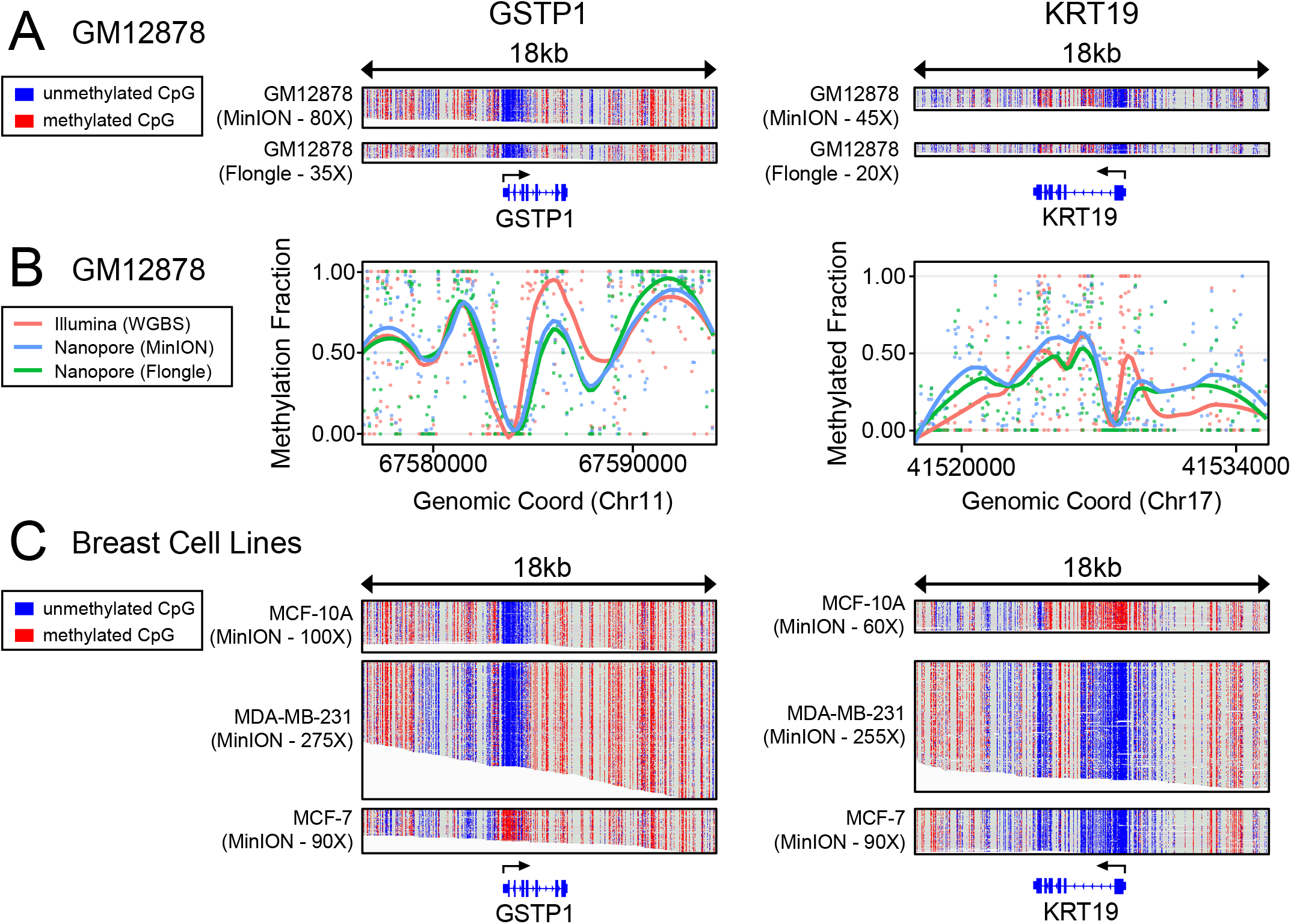
Methylation Studies and Comparison with Expression Data. Data shown for *GSTP1* & *KRT19*; 3 additional genes in Supplemental Figure S1. (A) Read-level methylation plots using IGV for GM12878 cell line at the promoter and gene body for GSTP1 and KRT19 (B) Comparison of Illumina WGBS data and Cas9-Nanopore methylation calls in the GM12878 cell line (C) Read-level methylation plots using IGV for the three breast cell lines (MCF-10A, MDA-MB-231, and MCF-7)

We use this technique to evaluate promoter methylation of the metabolism gene, glutathione S-transferase pi 1 (*GSTP1*), between breast cell lines. *GSTP1* is known to demonstrate promoter hypermethylation and transcriptional down-regulation in aggressive ER-positive breast cancer^16^. Correspondingly, we observe a dramatic increase in methylation at the *GTSP1* promoter in MCF-7: the only cell ER(+) breast cancer cell we examined (**Figure 2B**). Another example is the keratin family member gene: *KRT19*. *KRT19* expression is normally restricted to developing epithelial layers and not in mature mammary tissue^18^. *KRT19* is known to be expressed in breast cancers with poor prognostic outlook^17^, and detection of *KRT19* has been used to demonstrate metastasis of breast cancer to lymph nodes and bone marrow^18^. We observe that the *KRT19* gene remains largely methylated in the non-tumorigenic MCF-10-A cell line, but the gene promoter becomes hypomethylated in both of the transformed cell lines, MCF-7 and MDA-MB-231 (**Figure 2B**). We compared gene expression levels using existing RNA-seq data^11^ from these breast cell lines (**Supplemental Figure S3**). This demonstrated that mRNA levels of these genes follow the canonical inverse correlation between gene activity and promoter methylation, supporting the notion that CpG methylation is indeed playing a regulatory role.

### Structural Variation

To validate the nCATS method for calling structural variations, we used available 10X genomics data from the Genome In a Bottle (GIAB) Consortium project^9^ to identify large deletions present in the GM12878 cell line. We selected three heterozygous deletions, two with sizes of ~70kb and one ~150kb. Guide RNAs were designed to flank the deletion breakpoints by 5kb, resulting in reads of ~10 kb on the deleted allele, and spanning the region between cut sites on the non-deleted allele. Using existing familial trio sequencing data on GM12878^8^, HapCUT2^20^ was used to phase the reads and compare read lengths and read counts achieved from each allele. Interestingly, we found that the allele containing the deletion, with the correspondingly shorter distance between the cut sites, demonstrated a dramatically higher total number of reads (**Figure 3A**). This reflects a bias against achieving reads >50kb likely introduced during DNA purification or library preparation and delivery to the pore. To confirm this size-bias, we performed similar parental-allele segregation on sites without SVs and did not observe bias towards either parental allele (**Supplementary Figure S4**). The alignment data was passed to the Sniffles variant caller^21^ which identified all 3 of the deletions within 10nt from the annotated breakpoints in existing GIAB data (**Supplementary Table S2**). We adjusted Sniffles parameters to call SVs as heterozygous if an allele was supported by even a very low amount (0.1%) of reads, as the imbalance of reads from the two alleles caused the software to initially identify these deletions as homozygous (see methods).

**Figure 3:**
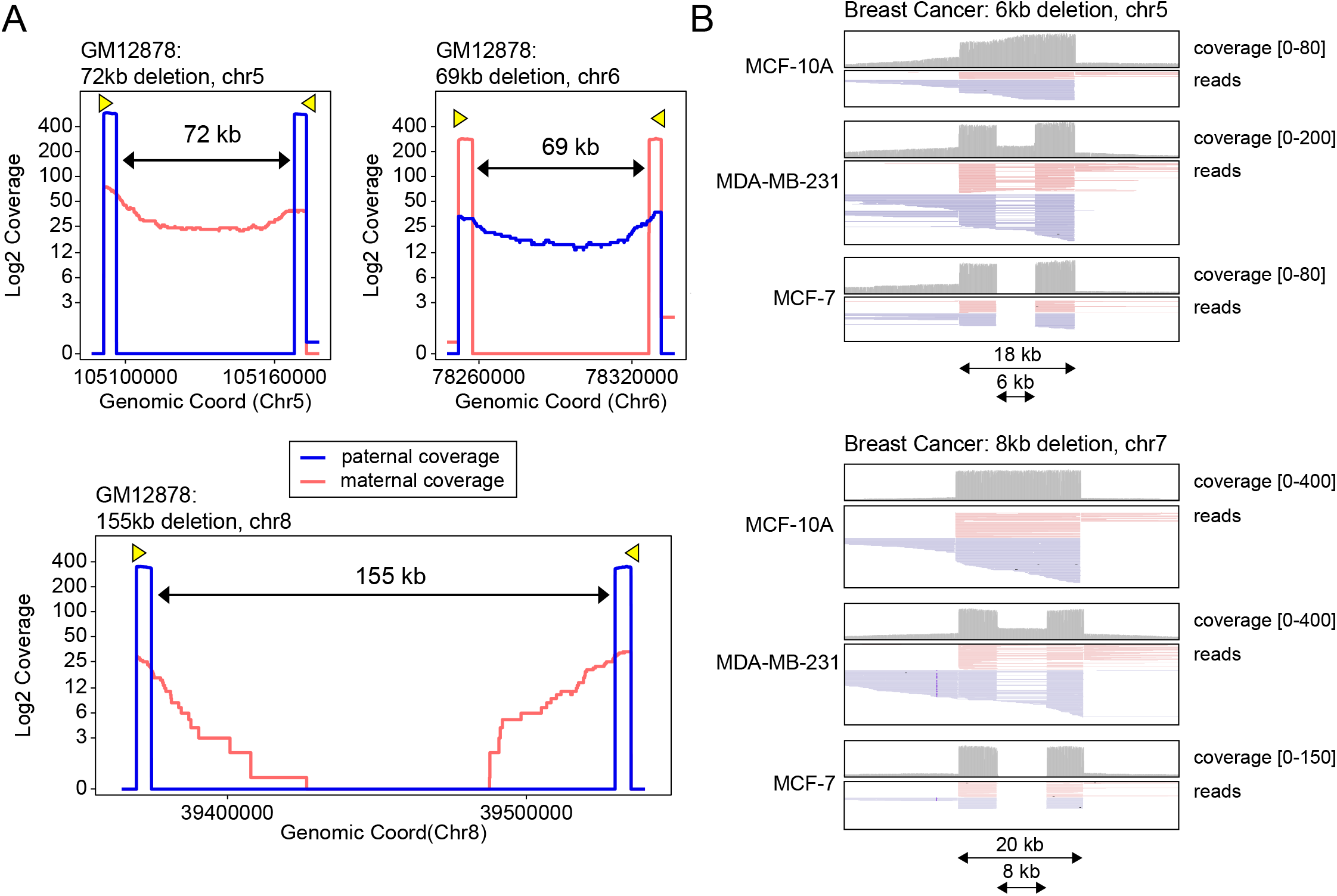
Structural Variation Studies. (A) Coverage data from three deletions in the GM12878 lymphoblast cell line, showing increased coverage from the deleted (shorter) allele. Reads were segregated into parental allele of origin using the haplotype-aware tool HapCUT217 with reference data from familial trio sequencing^9^. Yellow triangles show Cas9 cut site and guideRNA direction (B) IGV plots showing reads and coverage at two deletions present in the MDA-MB-231 and MCF-7 cell line and absent in the MCF-10A cell line.

Using candidate deletions from whole genome nanopore sequencing data previously published from our lab^10^, we selected two deletions present in the MDA-MB-231 and MCF-7 breast cancer lines and absent in the MCF10A cell line. Plotting reads around these loci in IGV showed both of these loci have heterozygous deletions in MDA-MB-231 and homozygous deletions in MCF-7 (**Figure 3B**). The SV caller Sniffles identified both of the suspected ~7kb deletions from alignment data calling the deletions as heterozygous in MDA-MB-231 and homozygous in the MCF-7 cell line (**Supplementary Table S3**). We also performed methylation studies on these regions but did not note any difference in methylation patterns between the deleted and not-deleted allele (**Supplementary Figure S5**). Together, this demonstrates how the high coverage nanopore data achieved through the nCATS method enhances our ability to evaluate and examine structural variants and can be combined with methylation calls to study CpG methylation patterns at deletion breakpoints.

We added the BRCA1 gene to our initial flongle flow cell testing because of the well-documented association of this gene with familial breast cancer^22^. BRCA1 was especially attractive as a target because of the abundance of hard to map repetitive Alu elements in the region^23^. To capture the entire BRCA1 gene, distance between flanking guideRNAs was 84kb, larger than areas we generally target. We only found a single read which spanned the entirety of the BRCA1 locus, with a regional coverage of 3X (**Supplementary Figure S6**). Further optimization is needed for >80kb nCATS to better resolve complex repetitive regions like this one.

### Single Nucleotide Variant Detection

Nanopore sequencing still has intrinsically high error rates (~10%) due to the inability of the basecaller to distinguish between some k-mers and the difficulty in discriminating signal events in repetitive regions (e.g. homopolymers). We explored how the increased coverage data from the nCATS protocol would affect the ability to call variants from nanopolish data. We compared the variant-calling performance of the samtools package^24^, which uses only alignment data, with the nanopolish variant caller^7^, which uses raw electrical data as well as alignment data. For initial validation, we once again turned to the GM12878 cell line, using the well-annotated platinum genome dataset as ground truth for SNVs^8^. Applying variant calling analysis to the 8 loci without deletions, we called SNVs over a total enriched area of 140kb which has 176 annotated SNVs. Following the most recent update to the basecaller (Guppy V3.0.3), we found that both samtools and nanopolish both performed well at calling single base substitutions (**Table 2**). Using data from the MinION run, samtools correctly identified 170 SNVs with 12 false positives on the default settings (**Supp Table S4A**; Sensitivity: 0.97, Positive predictive value (PPV): 0.93) and nanopolish identified 169 (**Supp Table S4B**; Sensitivity: 0.96, PPV: 0.91). In the lower coverage data from the Flongle run, samtools was only able to call 138 SNVs (**Supp Table S5A**; Sensitivity: 0.78, PPV: 93), while nanopolish performed better with 160 variants correctly found (**Supp Table S5B**; Sensitivity: 0.91, PPV: 0.89). Nanopolish is able to perform better with less data because of its ability to add the electrical data while interrogating sequence information for identifying SNVs.

**Table 2:**
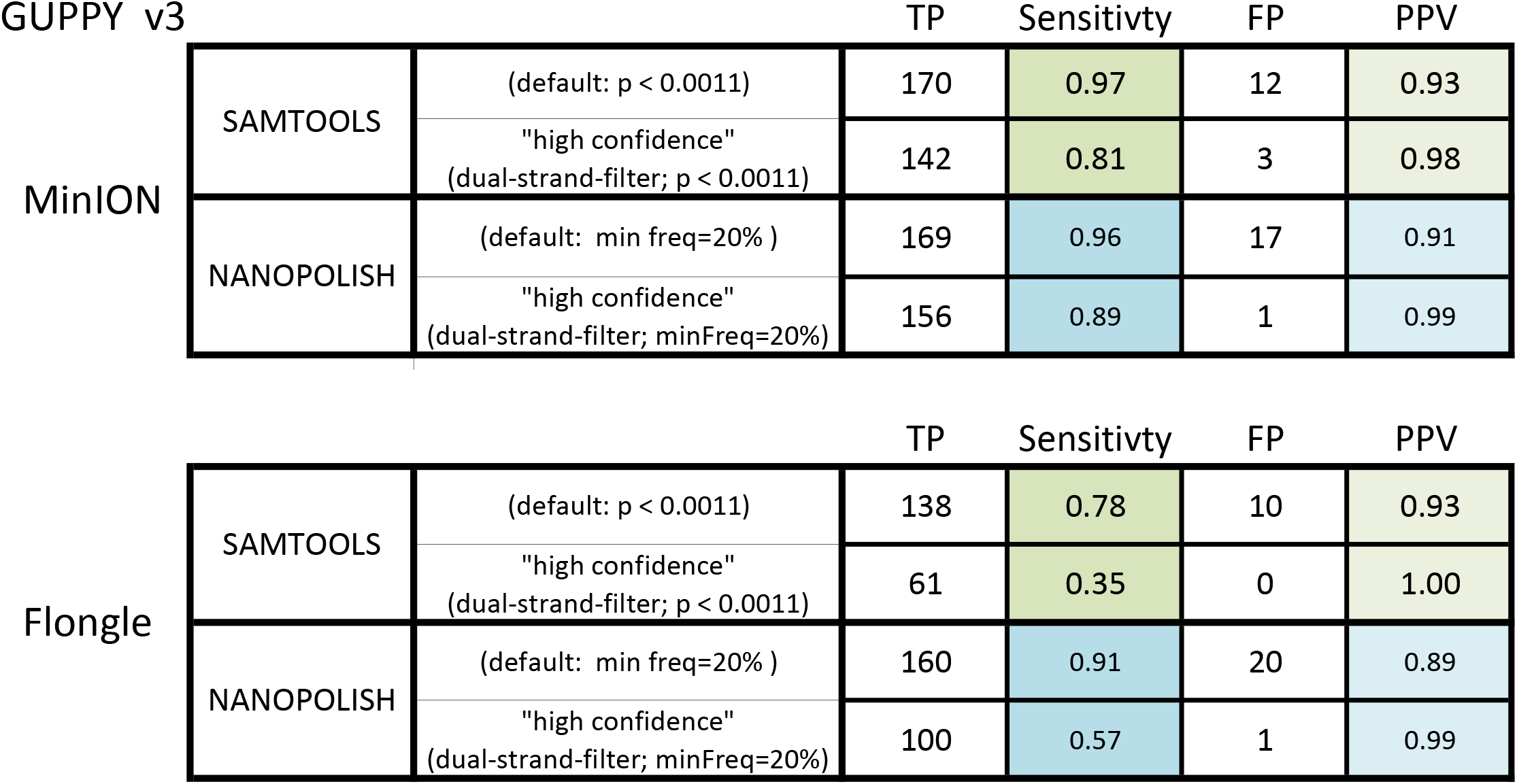
176 annotated variants are present in the 140kb (total) being queried across all sites. Comparing variant calling performance with samtools and nanopolish software tools, both on raw data as well as requiring variants to be supported by data from both strands (dual-strand-filter). Sensitivity = (True positives result / All true positives) Positive Predictive Value = (True positive result / All positive results)

MinION
|  |  | TP | Sensitivity | FP | PPV |
| --- | --- | --- | --- | --- | --- |
| SAMTOOLS | (default: $p < 0.0011$) | 170 | 0.97 | 12 | 0.93 |
| | "high confidence"<br>(dual-strand-filter; $p < 0.0011$) | 142 | 0.81 | 3 | 0.98 |
| NANOPOLISH | (default: min freq=20% ) | 169 | 0.96 | 17 | 0.91 |
|  | "high confidence"<br>(dual-strand-filter; minFreq=20%) | 156 | 0.89 | 1 | 0.99 |

**Table 2:** 176 annotated variants are present in the 140kb (total) being queried across all sites.
|  |  | TP | Sensitivity | FP | PPV |
| --- | --- | --- | --- | --- | --- |
| SAMTOOLS | (default: p < 0.0011) | 138 | 0.78 | 10 | 0.93 |
|  | "high confidence"<br>(dual-strand-filter; p < 0.0011) | 61 | 0.35 | 0 | 1.00 |
| NANOPOLISH | (default: min freq=20% ) | 160 | 0.91 | 20 | 0.89 |
|  | "high confidence"<br>(dual-strand-filter; minFreq=20%) | 100 | 0.57 | 1 | 0.99 |

The relatively high rate of false positive variants (5-10%) makes it difficult to discern true variants from those that are the result of sequencing errors, hindering the use of this method for *de novo* SNV identification. On inspection, we noted many false positives to occur from errors on only one strand (**Supplementary Figure S7**). Presumably, the basecaller was having systematic issues with correct identification of nucleotides on one strand but not the reverse complement. To reduce the strand-specific false positives, we implemented a filter requiring identified variants to be supported by reads from both strands. This filter did cause a modest decrease in sensitivity (**Table 2**) but eliminated all false positives except one in the nanopolish SNV calls (a highly thymidine-dense homopolymer region, **Supplementary Figure 8**). A visual representation of variants detected by the dual-strand filter at the *TP53* locus from both MinION and Flongle data is shown in **Figure 4**, for which raw nanopore reads were error-corrected with the ‘phase-reads’ module of nanopolish and the *de novo* identified variants.

**Figure 4:**
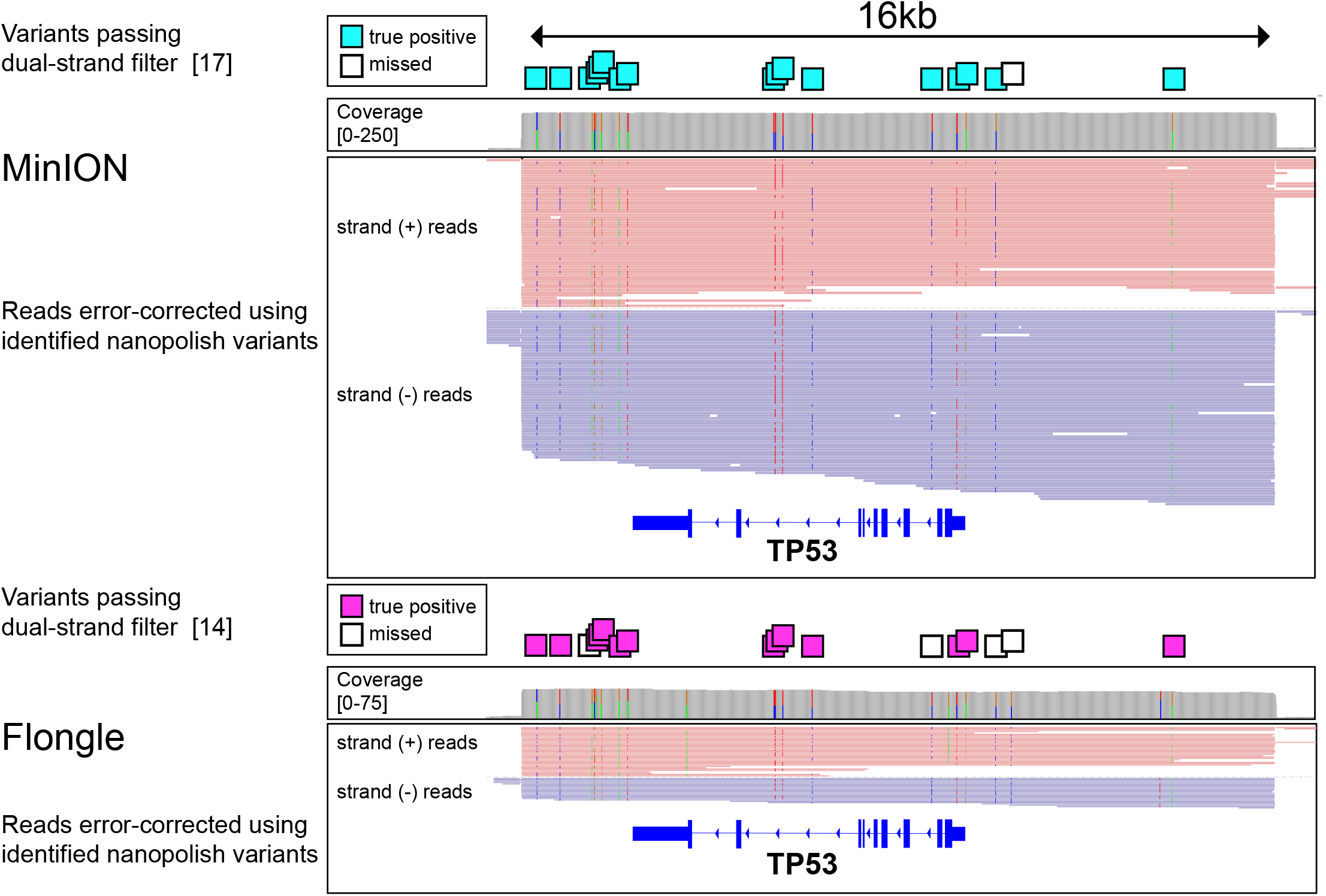
SNV detection IGV plot of the captured region around the TP53 gene in GM12878. Reads were error-corrected using the ‘phase-reads’ nanopolish module and all variants called by nanopolish for both the MinION data (top) and flongle data (bottom). Boxes above the region show SNVs in the platinum genome reference^9^, colored boxes represent detected variants passing the dual-strand filter. Three false positive variants are visible in the flongle data IGV plots; no false positives were present in the MinION data in this region.

Applying this variant calling pipeline to the MDA-MB-231 cell line, we selected 3 annotated single nucleotide variants^25^ in genes causally implicated in cancer (*BRAF*, *KRAS*, and *TP53*). Across these 3 regions, nanopolish called 42 SNVs in MDA-MB-231 from our MinION data set. For comparison, the GM12878 cell line has 40 annotated SNVs in these three regions, but in GM12878 variants are restricted to introns and UTRs, unlike in MDA-MB-231 where mutations exist in exons and affect protein coding. The known mutations in BRAF, KRAS, and TP53 were called with the correct ploidy using nanopolish (**Supplementary Table 4**). The mutations in BRAF and TP53 both passed the dual-strand filter, but the variant in KRAS was only detected on the positive strand. Taken together, the nCATS enrichment method provides a strategy for identification of single nucleotide variants *de novo* from nanopore signal, and can be used to phase and visualize known variants.

## Discussion

We have described a new method that can be rapidly adapted to evaluate a number of genomic sites simultaneously with high coverage nanopore sequencing data. Because of the low cost to entry and small footprint of the instrument, this assay has the potential to be widely utilized as a tool for evaluating DNA methylation, structural variation, and even small mutations. We show that single nucleotide variants in regions of interest can be queried with the nCATS protocol, although there are persisting limitations of nanopore variant calling at present, as evinced by the few SNVs not detected by this approach. As basecaller algorithms continue to improve we anticipate even higher future performance of this tool for the surveillance and identification of mutations. We also highlight the use of the nCATS method to detect and reassess structural variants. It is only recently, with the advent of long-read sequencing that the great diversity of structural variation in human genomes has been appreciated^26,27^, and this method provides a dynamic tool to evaluate genomic rearrangements, which contribute to cancer pathogenesis^28^. We show that the assay can be used to evaluate the ploidy of SVs and segregate reads into their parental alleles. Importantly, because nanopore sequencing interrogates the DNA strand rather than sequencing “by-synthesis”, we can *simultaneously* profile methylation in these loci, providing biological as well as diagnostic insight into the epigenome, which is commonly disrupted in human neoplasia^29^. By positioning guide RNAs adjacent to low complexity regions, this technique can also evaluate DNA methylation in regions of poor bisulfite sequencing mappability^30^ while avoiding the high costs associated with whole genome bisulfite sequencing. Thus, this assay represents a fast and inexpensive way to assess clinically relevant genes to detect genomic or epigenomic variation.

The strategies presented here also offer tools for additional applications including locating gene insertions from intentional genetic engineering or viral integration, evaluating tandem repeats present in pathological regions, and generating long-read data for building genome assembly scaffolds. By providing a targeted approach that can be rapidly applied to studying DNA at numerous regions of interest, this method helps to expand the applications of nanopore sequencing as an inexpensive alternative for generating high coverage targeted data while maintaining the advantages of long-read sequencing through directly probing the native cellular DNA.

## METHODS

### Cell culture and DNA prep

Cell lines were obtained from ATCC and cultured according to standard protocols. DNA was extracted using the MasterPure kit (Lucigen, MC85200), and stored at 4C until use. DNA was quantified using the Qubit fluorometer (Thermo) immediately before performing the assay.

### Guide RNA design

Guide RNAs were assembled as a duplex from synthetic crRNAs (IDT, custom designed) and tracrRNAs (IDT, 1072532). Sequences are provided in **Supplementary Table S1**. The crRNAs were designed using IDT’s design tool and selected for the highest predicted on-target performance with minimal off-target activity. The gRNA duplex was designed to introduce cuts on complementary strands flanking the region of interest. For methylation studies and SNV studies, the target size between gRNAs was 20-30 kb; for deletions, the gRNAs were designed to flank the suspected breakpoints by ~5kb.

### Ribonucleoprotein Complex Assembly

Prior to guide RNA assembly, all crRNAs were pooled into an equimolar mix, with a total concentration of 100uM. The crRNA mix and tracrRNA were then combined such that the tracrRNA concentration and total crRNA concentration were both 10uM. The gRNA duplexes were formed by denaturation for 5 minutes at 95C, then allowed to cool to room temp for 5 minutes on a benchtop. Ribonucleoprotein complexes (RNPs) were constructed by combining 10pmol of gRNA duplexes with 10pmol of HiFi Cas9 Nuclease V3 (IDT, 1081060) in 1X CutSmart Buffer (NEB, B7204) at a final volume of 30uL (conc: 333nM), incubated 20 minutes at room temperature, then stored at 4C until use, up to 2 days.

### Cas9 Cleavage and Library Prep

3ug of input DNA was resuspended in 30uL of 1X CutSmart buffer (NEB, B7204), and dephosphorylated with 3uL of Quick CIP enzyme (NEB, M0508) for 10 min at 37C, followed by heating for 2 minutes at 80C for CIP enzyme inactivation. After allowing the sample to return to room temp, 10uL of the pre-assembled 333nM Cas9/gRNA complex was added to the sample. In the same tube, 1uL of 10mM dATP (Zymo, D1005) and 1uL of Taq DNA polymerase (NEB, M0267) were added for A-tailing of DNA ends. The sample was then incubated at 37C for 20min for Cas9 cleavage followed by 5 minutes at 72C for A-tailing. Sequencing adaptors and ligation buffer from the Oxford Nanopore Ligation Sequencing Kit (ONT, LSK109) were ligated to DNA ends using Quick Ligase (NEB, M2200) for 10 min at room temp. The sample was cleaned up using 0.3X Ampure XP beads (Beckman Coulter, A63881), washing twice on a magnetic rack with the long-fragment buffer (ONT, LSK109) before eluting in 15uL of elution buffer (ONT, LSK109). Sequencing libraries were prepared by adding the following to the eluate: 25uL sequencing buffer (ONT, LSK109), 9.5uL loading beads (ONT, LSK109), and 0.5uL sequencing tether (ONT, LSK109).

### Sequencing

Each sample was run on a 9.4.1 version flow cell using the GridION sequencer. Initial flow cell priming was performed with 800uL flush buffer (ONT, LSK109), and allowed 5 minutes to equilibrate. The flow cells were then primed with 200uL of a 1-to-2 dilution of sequencing buffer (ONT, LSK109) prior to loading sequencing libraries.

### Analysis

Basecalling was performed using the GUPPY algorithm (Version 3.0.3) to generate FASTQ sequencing reads from electrical data. Reads were aligned to the human reference genome (Hg38) using either NGMLR^21^ (for SV calling) or Minimap2^31^. Per-nucleotide coverage was determined using samtools, and clustered using the ‘bincov’ script of the SURVIVOR software package^32^.

CpG methylation calling on nanopore data was performed using nanopolish^7^. Methylation calling on existing WGBS data of GM12878 (GEO: GSE86765)^19^ was performed using the bismark software tool^33^. RNA-seq data of MCF-10A, MCF-7, and MDA-MB-231 were downloaded from GEO (Accession: GSE75168) in the form of RNA counts.

Deletions were called using the structural variant caller Sniffles^21^, set to find deletions with a minimum size of 100bp. Because of the allelic size bias on the very large deletions in GM12878, the ploidy was initially incorrectly called as homozygous. To correct this, we used the option “--min_homo_af” set to 99.9, which ensured a deletion was called as heterozygous if reads supporting the reference were present at a minimum threshold of one in one thousand.

Segregation of GM12878 reads into parental alleles was performed using HapCUT2^20^ with existing phased variant information from sequencing of familial trios^8^. For assignment into parental allele, we required at least 75% of variants per read to agree on allele of origin. If variants were not detected or if the variants within a read disagreed on the parental allele it was excluded from allelic analysis.

*De novo* variant calling was performed using nanopolish^7^ or samtools^24^. For calling true positives, we used the platinum genome dataset^8^, filtered to contain only SNVs. For calling specificity, the total number of tests was taken as all regions queried with the most permissive nanopolish settings (minimum freq: 10%, minimum quality score: 0).

### Accession

Cas9-Enrichment data are available at NCBI Bioproject ID PRJNA531320 (https://www.ncbi.nlm.nih.gov/bioproject/PRJNA531320). Code used in data analysis is available at https://github.com/timplab/Cas9Enrichment

