## Supplemental Material for "Targeted Nanopore Sequencing with Cas9 for studies of methylation, structural variants, and mutations"

**Supplemental Materials**

**Supplemental Figures**

Fig S1: Methylation read-level plots, and methylation line plots for additional 3 genes
Fig S2: Per CpG methylation comparison between nanopore calls and Illumina calls
Fig S3: Normalized RNA-seq read counts for 5 genes in breast cell lines
Fig S4: Figure showing allelic bias not present at other loci without deletions
Fig S5: Methylation at 2 SVs in breast cell lines
Fig S6: Reads for the BRCA1 gene from the Flongle sequencing run of GM12878
Fig S7: False positive variants and true positive variants demonstrating the impetus for implementing a dual-strand filter
Fig S8: Sole false positive variant found by nanopolish that passes dual-strand filter

**Supplemental Tables**

(Supp Tables 1, 4, 5, 6 as Excel sheets; Supp Tables 2, 3 in this document)

Table S1: GuideRNA targets, sequences, and loci
Table S2: Sniffles Calls of large SVs in GM12878
Table S3: Sniffles Calls SVs in 3 breast cell lines
Table S4: SNVs called in GM12878 MinION data (8 loci)
Table S5: SNVs called in GM12878 Flongle data (8 loci)
Table S6: SNVs called in MDA-MB-231 MinION data (3 loci)

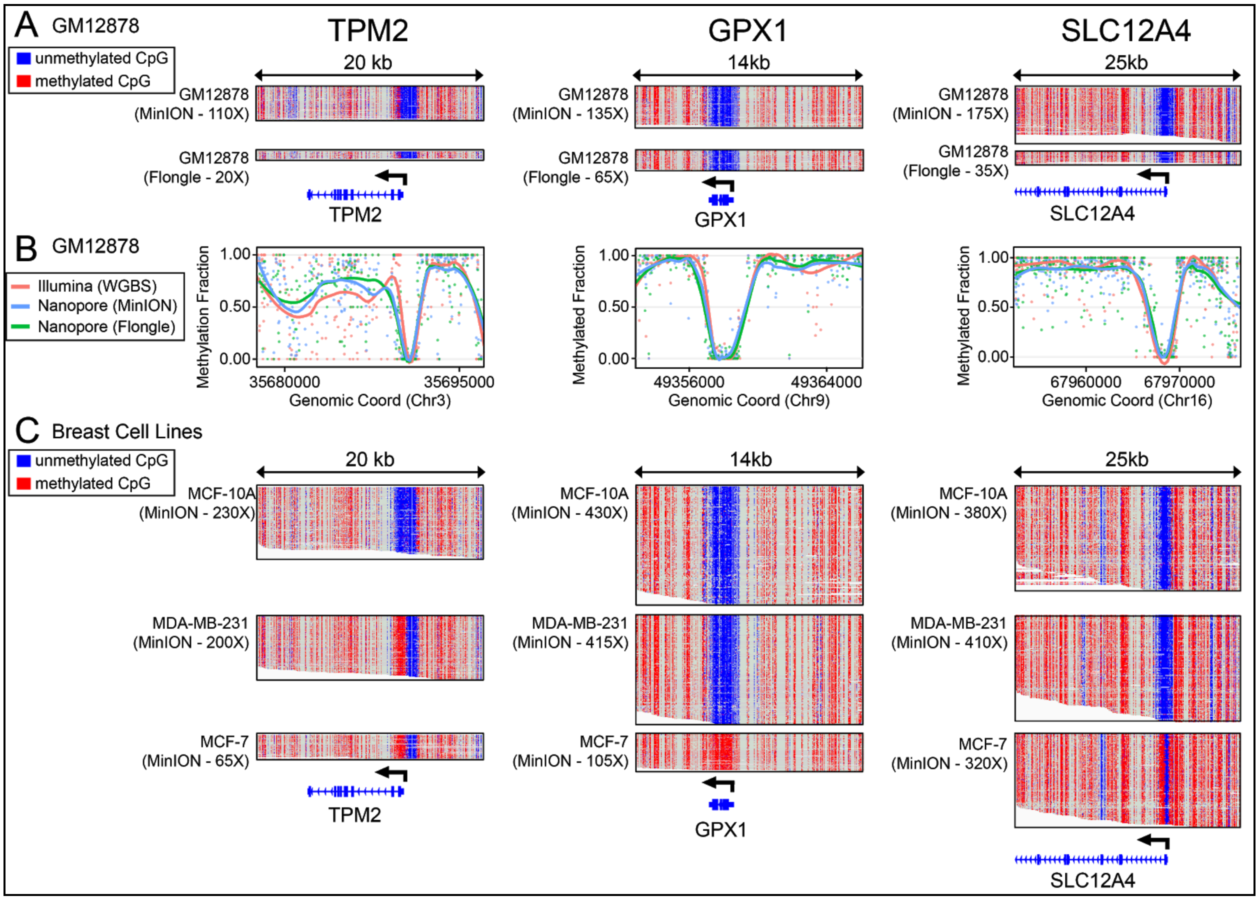


**Figure S1:** (A) Read-level methylation plots using IGV for GM12878 cell line at the promoter and gene body for *TPM2*, *GPX1*, and *SLC12A4*. (B) Comparison of Illumina WGBS data and Cas9-Nanopore data in the GM12878 cell line. (C) Read-level methylation plots using IGV for the three breast cell lines (MCF-10A, MDA-MB-231, and MCF-7)

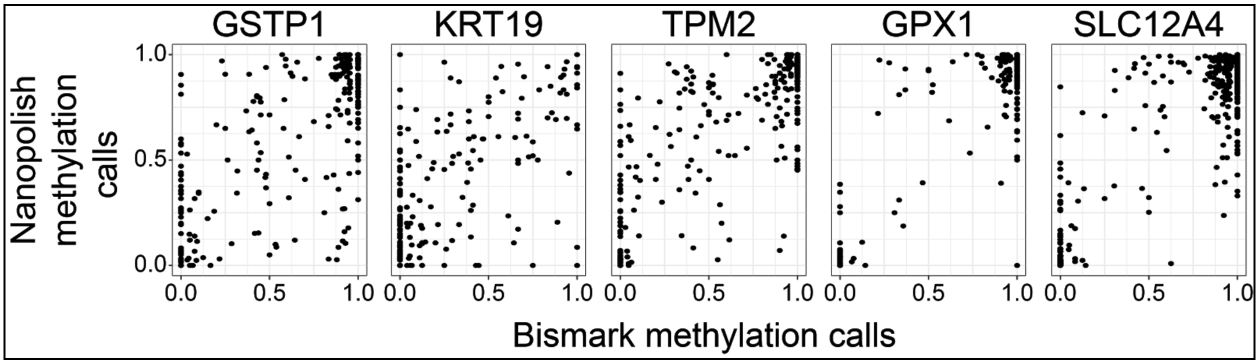


**Figure S2:** Comparing methylation calls made by bismark (WGBS Illumina data: GEO: GSE86765) and nanopolish (Cas9-targeted nanopore data) at all CpG in the targeted regions. $R^2$ = 0.82 across all 5 sites.

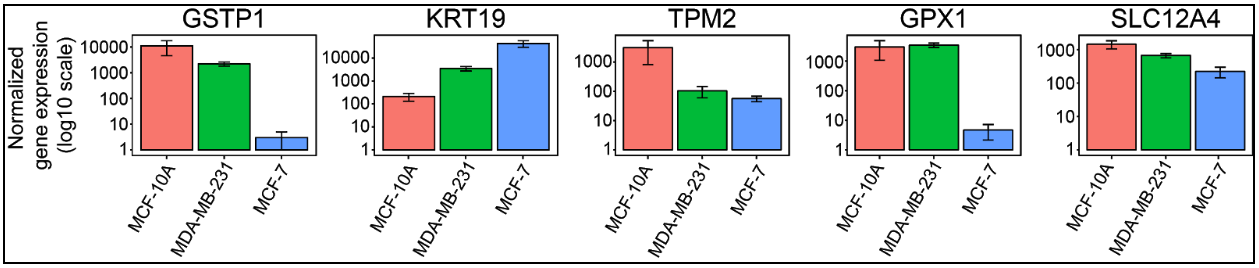


**Figure S3:** Normalized expression data for three breast cell lines from existing RNA-seq data (GEO: GSE75168).

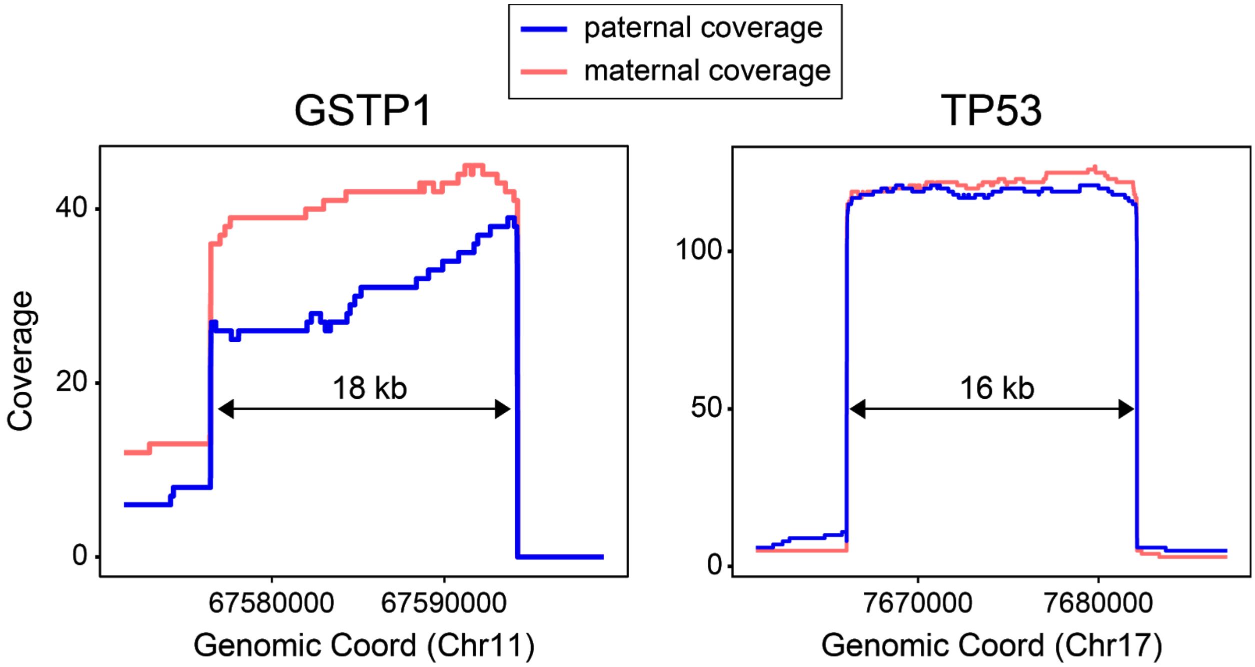


**Figure S4:** Comparing paternal and maternal coverage at two sites in GM12878 with no heterozygous SVs between guideRNAs. Unlike at the sites of large heterozygous deletions, we do not see a dramatic bias towards either parental allele.

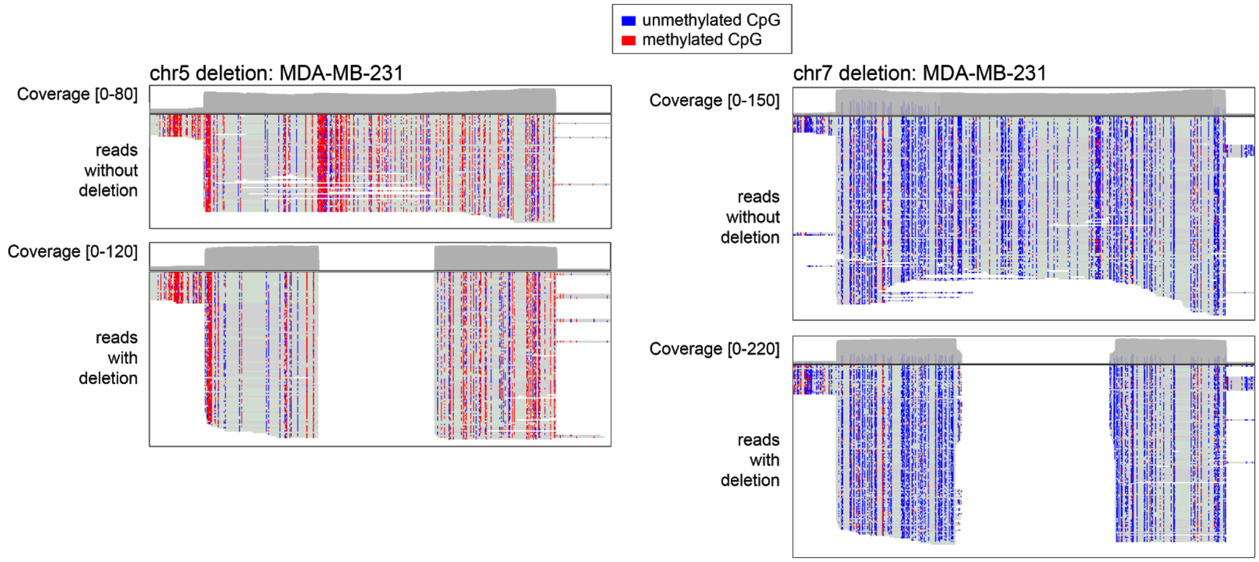


**Figure S5:** Comparing methylation patterns at heterozygous deletions on chr5 and chr7. No difference in methylation was observed between in reads with the deletion versus reads without the deletion.

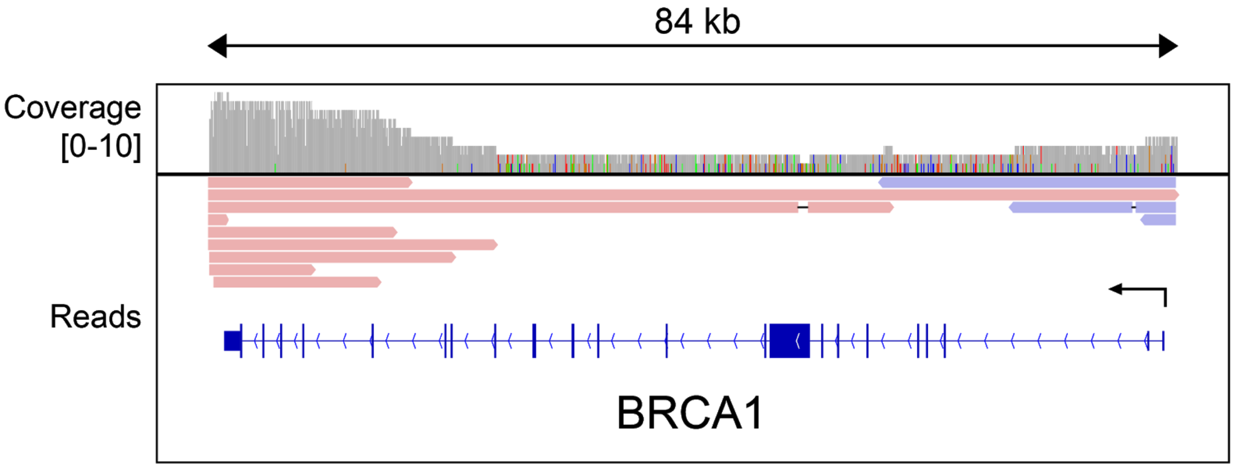


**Figure S6:** The few reads resulting from trying to target the larger BRCA1 gene in GM12878 cell line using the flongle flow cell. We found only one read spanning the entirety of the gene

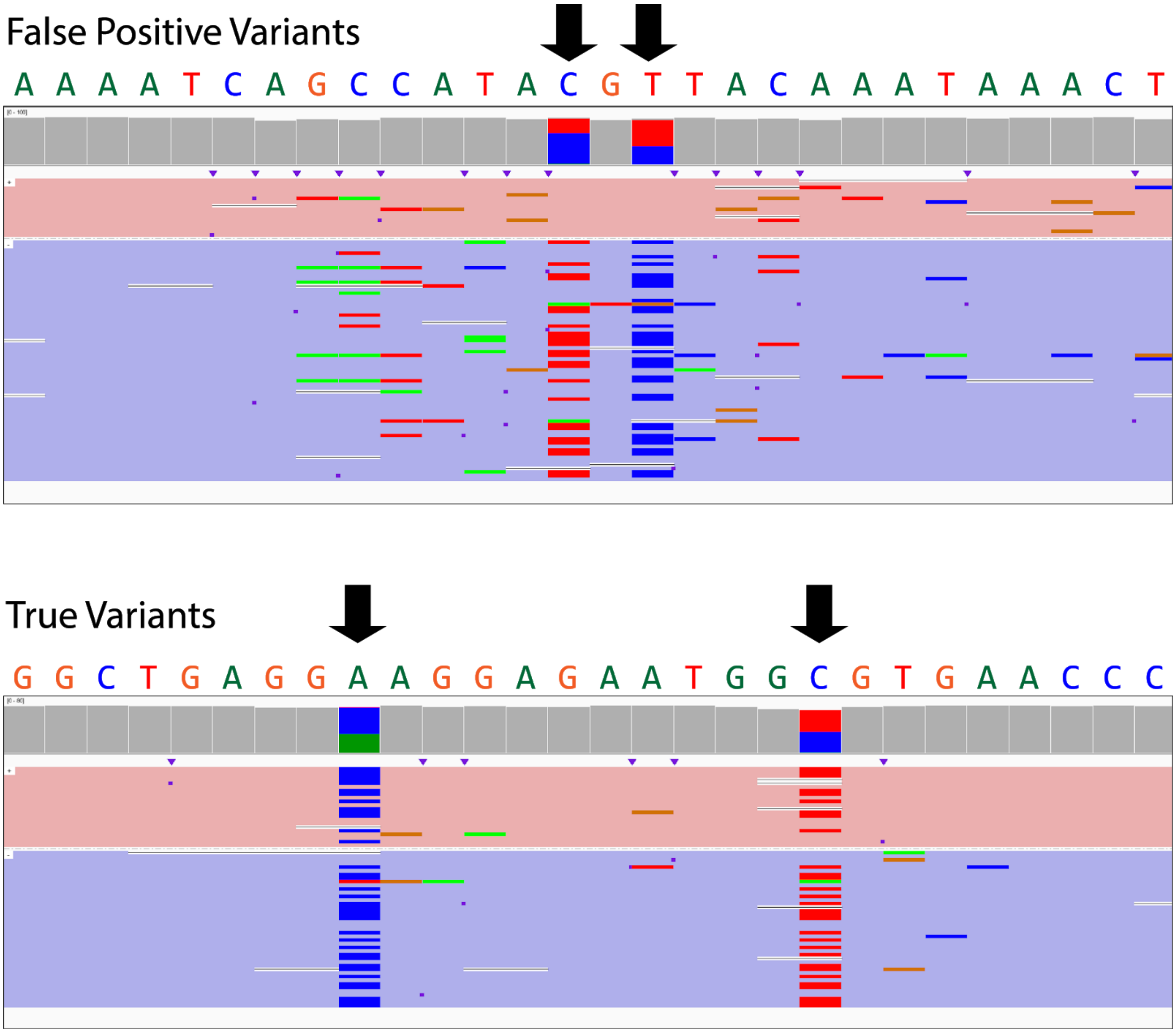


**Figure S7:** Example of two false positive variants resulting from an error on only one strand, and two real variants which are supported by data on both strands

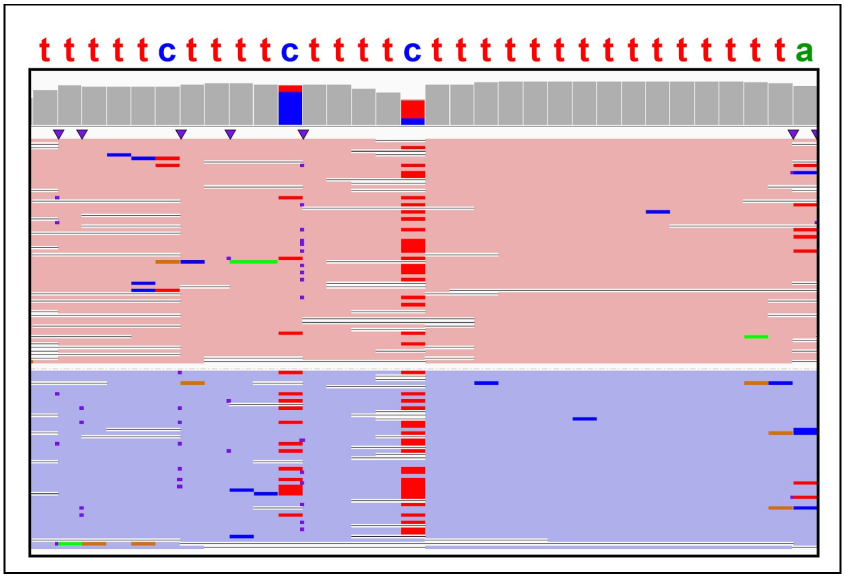


**Figure S8**: The single false positive variant found by nanopolish that passes dual-strand filter, present in a highly thymidine-dense region

SUPPLEMENTARY TABLES

**Table S1** (Excel Spreadsheet): A) GuideRNA sequences and target sites for the 10 + BRCA1 targeted regions for methylation, SV and SNV interrogation. B) GuideRNA sequences and target sites for the 3 additional large deletions in GM12878. All target locations are given in hg38 coordinates

GM12878 - Large Deletions

REFERENCE

Reference Calls (GIAB, 10X genomics LongRanger2.1 data)

| chromosome | start site | size | GT |
|---|---|---|---|
| chr5 | 105096420 | 71545 | het |
| chr6 | 78257486 | 69265 | het |
| chr8 | 39374563 | 155140 | het |

CALLED

Sniffles Calls (min size: 100bp; min support: 50 reads)

| chromosome | start site | size | GT* |
|---|---|---|---|
| chr5 | 105096412 | 71560 | het |
| chr6 | 78257496 | 69263 | het |
| chr8 | 39374556 | 155153 | het |

**Table S2:** (Left) Reference calls from LongRanger 2.1 analysis of 10X genomics data from the GIAB consortium(Zook et al. 2016). (Right) SNIFFLES SV calls in GM12878. het = heterozygous

GT* - the genotype of these deletions in GM12878 was only correctly called by adjusting settings permitting the reference allele to present at a lower threshold (see methods)

Breast Cancer Cell Lines - Sniffles Calls
(min size: 100bp; min support: 50 reads)

**chr5 deletion**
region between guideRNAs: 58378092-58396781

| cell line | start site | size | GT |
|---|---|---|---|
| MCF-10A | [no SV called] | [] | [] |
| MDA-MB-231 | 58384160 | 6107 | het |
| MCF-7 | 58384160 | 6110 | homo |

**chr7 deletion**
region between guideRNAs: 154593967-154613978

| cell line | start site | size | GT |
|---|---|---|---|
| MCF-10A | [no SV called] | [] | [] |
| MDA-MB-231 | 154600417 | 7569 | het |
| MCF-7 | 154600417 | 7569 | homo |

**Table S3:** Sniffles calls from enrichment data in the 3 breast cancer cell lines. For both deletions the ploidy was called as heterozygous in MDA-MB-231 and homozygous for MCF-7, in line for what we observed in IGV plots of the reads. het = heterozygous ; homo = homozygous

**Table S4** (Excel Spreadsheet): MinION data - Single nucleotide variants in GM12878 cell line identified *de novo* from nanopore signal at eight loci (*TP53, BRAF, KRAS, GSTP1, KRT19, TPM2, GPX1, SLC12A4*).
(S4A) Samtools; (S4B) Samtools w/ dual-strand-filter;
(S4C) Nanopolish; (S4D) Nanopolish w/ dual-strand filter

**Table S5** (Excel Spreadsheet): Flongle data - Single nucleotide variants in GM12878 cell line identified *de novo* from nanopore signal at eight loci (*TP53, BRAF, KRAS, GSTP1, KRT19, TPM2, GPX1, SLC12A4*).
(S5A) Samtools; (S5B) Samtools w/ dual-strand-filter;
(S5C) Nanopolish; (S5D) Nanopolish w/ dual-strand filter

**Table S6** (Excel Spreadsheet): Single nucleotide variants in MDA-MB-231 cell line identified *de novo* from nanopore data at three loci (*TP53, BRAF, KRAS*).
(S6A) Nanopolish; (S6B) Nanopolish w/ dual-strand filter
